# Optogenetic Isolation of the Amacrine Cell-OFF Bipolar Cell Synapse Shows Selective D1R Control of Glycinergic Transmission

**DOI:** 10.64898/2025.11.29.691316

**Authors:** Timothy D. Maley, Andrea J. Wellington, Erika D. Eggers

## Abstract

**Purpose:** Light evoked inhibition of OFF cone bipolar cells (OFF BCs) is modulated both by background light levels and the action of dopamine through the dopamine D1 receptor (R). Since D1Rs are localized throughout the mouse retina, it is not known where in the light signaling pathway dopamine is modulating signals to OFF BCs. Here we tested a technique that allowed for the isolation of the amacrine cell (AC) to OFF BC circuit to determine if there are local D1R-induced changes in inhibition from presynaptic ACs onto OFF BCs.

**Methods:** We utilized the B6.Cg-Tg(Slc32al-COP4*H134R/EYFP) mouse line that expresses ChR2 in all the inhibitory cells in the retina. ACs expressing ChR2 were directly activated by light, while blocking photoreceptor mediated inputs. Inhibitory synaptic currents or ChR2-evoked excitatory currents were recorded using whole-cell patch clamp electrophysiology.

**Results:** Optogenetically activated inhibition from ACs elicited inhibitory currents that had slow kinetics similar to light-evoked inhibition, suggesting that they are using native synaptic mechanisms. We recorded optogenetically activated inputs to OFF BCs from pharmacologically isolated GABAergic and glycinergic input while photoreceptor inputs were blocked. D1R activation reduced glycinergic inputs to OFF BCs while leaving GABAergic inputs intact.

**Conclusions:** D1R modulation of isolated optogenetic activation of AC-OFF BC synapses showed similar changes to previous experiments with light-evoked inhibition to OFF BCs. Together, this suggests optogenetic activation of ACs can be used to understand how dopamine differentially shapes inhibitory changes in the inner retina.

## Introduction

The 24-hour day/night cycle creates an environment where many visual species must continually adapt to large changes in background luminance. Light levels can vary by more than 10^12^ over the course of a day. To maintain visual acuity under these wide ranges, retinas use parallel pathways. Rod photoreceptors are sensitive under lower light levels while cones are sensitive under brighter conditions ^1^. In addition, the photoreceptors themselves modulate sensitivity to background light ^2^.

Downstream of the photoreceptors, light adaptation of the bipolar cells (BCs) has also been shown to modify excitability independent of changes in photoreceptor sensitivity. Previous work has shown both OFF cone BCs (OFF BCs) and rod BCs have reduced inhibitory inputs under light adapted conditions ^3–5^. Increasing light levels are also associated with increased dopamine release from the dopaminergic amacrine cells (ACs) ^6, 7^. Several studies have suggested dopamine agonists can mimic the light adapted reductions in inhibition seen in the inner retina and ganglion cells ^8–14^. Dopamine type 1 receptors (D1Rs) are the dominant dopamine receptors in the inner retina of the mouse and are located on the OFF and ON cone BCs as well as GABAergic and glycinergic ACs and several ganglion cell types ^15–17^. Together, this suggests that dopamine controls neuromodulation of inner retinal circuitry in response to increasing luminance.

Since D1Rs are expressed by horizontal cells and subtypes of cone BCs and ACs, D1R modulation of OFF BC inhibition could happen at many locations in the retina ^18, 19^. Inhibition to OFF BCs also requires activation of both photoreceptors, BCs and ACs by light which influences the properties of the inhibition. The goal of this study was to address this issue by directly activating presynaptic ACs using optogenetics while recording inhibitory inputs on OFF BCs. We found this was a reliable way to elicit inhibition independent of upstream photosensitive inputs from the outer retina or melanopsin expressing retinal ganglion cells or modulation of BC inputs to ACs. We found that direct activation of ACs produced unique modulation of D1R inhibitory inputs compared to previously recorded light-evoked and spontaneous currents on OFF BCs.

## Methods

### Retinal Slice Preparation

Retinas were isolated from 8-11 week old B6.Cg-Tg (Slc32al-COP4*H134R/EYFP)^20^ male and female mice (Jackson Laboratories, Bar Harbor, ME) that contain Channel rhodopsin 2 (ChR2) under the VGAT promoter in the inhibitory ACs and horizontal cells. VGAT is the vesicular transporter for GABA and glycine and so should be expressed in all inhibitory neurons. All procedures and experiments were performed under ambient room light. Eyes were enucleated after euthanasia with carbon dioxide. After enucleation, the cornea and lens were removed, and eyecups were bathed in chilled extracellular solution that was bubbled with 95% O_2_-5% CO_2_ (see whole-cell recordings) with 800 U/ml of hyaluronidase for 20 min to remove the remaining vitreous. The eyecups were then washed with extracellular solution and retinas were isolated. The periphery of the retinas was trimmed to produce a flat square, and the retinas were mounted to 0.45-µm nitrocellulose filter paper (Millipore, Billerica, MA). The mounted retinas were cut into 250-µm-thick slices on a custom hand chopper, and the slices were mounted with vacuum grease to glass cover slips and transferred to a dark box while bubbled with 95% O_2_-5% CO_2_. Animal protocols conformed with the ARVO statement for the use of Animals in Ophthalmic and Visual Research and were approved by the University of Arizona Institutional Animal Care and Use Committee.

### Whole-cell Recordings

Extracellular solutions for dissections and whole-cell recordings were bubbled with 95% O_2_-5% CO_2_ compressed gas and contained the following (in mM): 125.0 NaCl, 2.5 KCl, 1.0 MgCl2, 1.25 NaH2PO4, 20.0 glucose, 26.0 NaHCO3, and 2.0 CaCl2, 2.0 Na-Pyruvate, 4.0 Na L-Lactate, 0.5 L-Glutamine. Intracellular solution for recording currents contained (in mM): 120.0 CsOH, 120.0 gluconic acid, 1.0 MgCl2, 10.0 HEPES, 10.0 EGTA, 10.0 tetraethylammonium-Cl, 10.0 phosphocreatine-Na2, 4.0 Mg-ATP, 0.5 Na-GTP, and 50.0 µM Alexa Fluor 568 (Thermo Fisher Scientific, Eugene, OR) and was adjusted to pH 7.2 with CsOH. Final osmolarity was confirmed with a micro osmometer, Precision Systems (Natick, MA).

Photoreceptor inputs to other cells in the retina were pharmacologically blocked with CNQX (25 µM), APV (50 µM), ACET (1 μM), and L-AP4 (50 µM) that block AMPARs, NMDARs, KainateRs and mGluR6s, respectively. Inhibitory inputs were isolated by using gabazine (20 µM), TPMPA (50 µM) or strychnine (1 µM) to block combinations of GABA_A_Rs, GABA_C_Rs and glycineR, respectively. D1Rs were agonized with SKF-38393 (SKF, 20 µM; Tocris, Bristol, United Kingdom). Solutions were perfused into the recording chamber by gravity perfusion system at a rate of ∼1ml/min. Solution outflow was forced with a peristaltic pump from Fisher Scientific (Hampton, NH). All chemicals were obtained from Sigma-Aldrich (St. Louis, MO), unless noted.

Whole-cell voltage clamp recordings of optogenetically (o) light-evoked inhibitory post-synaptic currents (oIPSCs) and ChR2 mediated excitatory currents (oECs) were measured on OFF BCs and ACs. Retinal slices on glass coverslips were placed in a custom 3D printed chamber and heated to 33°C by a temperature controlled inline heater Warner Instruments (Holliston, MA). For each recording, at least 3 oIPSCs were recorded for each condition, and incubation in extracellular solution with drugs to block photoreceptor inputs (photoblock – PB), the addition of drugs to isolate receptors types to the PB cocktail and the addition of SKF-38393 to the PB+receptor isolation cocktail each occurred for 5 minutes prior to recordings. oIPSCs from OFF BCs and ACs were measured while clamping at 0 mV, the reversal potential for nonselective cation channels. oECs were measured while clamping at -60 mV, the reversal potential for Cl^-^channels. Series resistance was compensated for in all recordings. Electrodes were pulled from borosilicate glass (World Precision Instruments, Sarasota, FL) using a P97 Flaming/Brown puller (Sutter Instruments, Novato, CA) and had resistances of 6–10 MΩ. Liquid junction potentials of 20 mV, calculated with Clampex software (Molecular Devices, Sunnyvale, CA), were corrected for before recording. Responses were sampled at 10 kHz and filtered at 6 kHz with the four-pole 165-Bessel filter on a Multi-Clamp 700B patch-clamp amplifier (Molecular Devices).

Confirmation of OFF BC morphology ^21^ ^22^ was done at the end of each recording using an Intensilight fluorescence lamp and Digitalsight camera controlled by Elements software (Nikon Instruments, Tokyo, Japan).

ChR2 expressing cells were optogenetically activated by a 5 ms or 1 sec. full-field light stimulus projected through a 4x or 60x objective (λ = 470 nm). Stimulus intervals were 20 s to 1min. The light stimulus was generated with a Mightex Polygon 400 pattern illuminator (Pleasanton, CA). Whole field illumination patterns were controlled by percent LED intensity. LED intensity was calibrated to light intensity as photons·µm^−2^·s^−1^ and controlled through Polyscan 2 software by Mightex. Light intensities were calibrated with an S471 optometer (Gamma Scientific, San Diego, CA).

### Data analysis and statistics

Each light stimulus and recording were repeated 3 times. The average peak amplitude, decay time constant (t), time to first peak, frequency and charge transfer (Q) were measured for all evoked responses. Any cells with seals reduced below 1 GΩ or not displaying currents were excluded from the analysis. Q was calculated from light stimulus onset to when the trace returned to baseline. Time to first peak was measured from light stimulus onset to peak amplitude of the first current. Decay time constants (t), were calculated using a standard exponential fit using Clampex software (Molecular Devices).

Measurements in SKF were normalized to control response in either PB or PB+inhibitory receptor antagonists for each cell. T-tests were used to compare the responses before and after SKF treatment. Light dependent changes in AC oECs were analyzed using one-way ANOVA tests. Differences were considered significant if the p value < 0.05. GraphPad (Prism Software, Boston, MA) and SigmaPlot (Systat Software, San Jose, CA) were used for creation of graphs and statistical calculations. Error bars are SEM.

### Immunohistochemistry, Imaging and Analysis

Eyes from ChR2-EYFP+ (or ChR2-EYFP-for negative control) were enucleated, and eye cups (with cornea & lens removed) were fixed for 30 minutes in 4% paraformaldehyde. Eye cups were rinsed in Phosphate Buffered Saline (PBS), then retinas were dissected out. Retinas were embedded in 5% agarose and sliced at 60 µm on a vibratome. Retina slices were transferred to a 24-well dish and incubated for 1 hour in blocking solution (5% goat serum, 0.5% Triton X-100 in Phosphate Buffered Saline). Retina slices were incubated overnight at room temperature in blocking solution using these primary antibodies and dilutions: Mouse IgG1 anti-GFP 1:200 (DSHB-GFP-8H11); Mouse IgG2a anti-GAD65 1:1000 (DSHB GAD-6); Mouse IgG2a anti-GAD 67 1:1000 (Millipore MAB5406); Rabbit anti-GlyT1 1:300 (Cedarlane #SY 272103). Slices were washed 3x for 30 min each in PBS, then incubated with the following secondary antibodies with 1:1000 dilution in PBS for 2.5 hours: Anti-mouse IgG1-AlexaFluor647 (Invitrogen A21240); Anti-mouse IgG2a-AlexaFluor488 (Invitrogen A21131); Anti-rabbit AlexaFluor 546 (Invitrogen A10040). Nuclei were stained with DAPI (4′,6-Diamidine-2′-phenylindole dihydrochloride) 1:1000 (Invitrogen #D1306) during secondary antibody incubation. Slices were washed 3x for 30 min each in PBS, then mounted in ProLong Glass (Invitrogen #P369820).

Slides were imaged on a Zeiss LSM 880 inverted confocal microscope using a 40x objective (Plan-Apochromat 40x/1.3; University of Arizona - UA ORP Imaging Cores - Optical core facility, RRID:SCR_023355). Z-stacks were acquired using 0.38 µm slice intervals. Images were collected as 16-bit, 212.55 x 212.55 µm (1024 x1024 pixels). The following laser intensities were used: GAD65/67 488nm laser at 2%, GlyT1 561nm laser at 4%, GFP 633nm laser at 8%, DAPI 405nm laser at 0.5%.

## Results

To confirm localization of ChR2-EYFP to the inhibitory neurons in the B6.Cg-Tg (Slc32al-COP4*H134R/EYFP – VGAT-ChR2-EYFP) mouse line ^20, 23^, retinal slices were immunolabeled for EYP (GFP), nuclei (DAPI), Glutamic Acid Decarboxylase (GAD65/67, GABAergic neurons), Glycine Transporter Type-1 (GlyT1, Glycinergic neurons) (Fig. 1). VGAT-ChR2-YFP^-^ mice did not show EYFP fluorescence (Fig. 1A) but in VGAT-ChR2-EYFP^+^ mice EYFP was expressed in the outer plexiform layer (OPL), inner nuclear layer (INL), and inner plexiform layer (IPL) (Fig. 1B). There was significant overlap of EYFP with GlyT1 (Fig. 1C-D) and GAD65/67 (Fig. 1E-F), showing expression in ACs. Rod BCs labeled with PKCalpha did not express EYFP (not shown).

**Figure 1:**
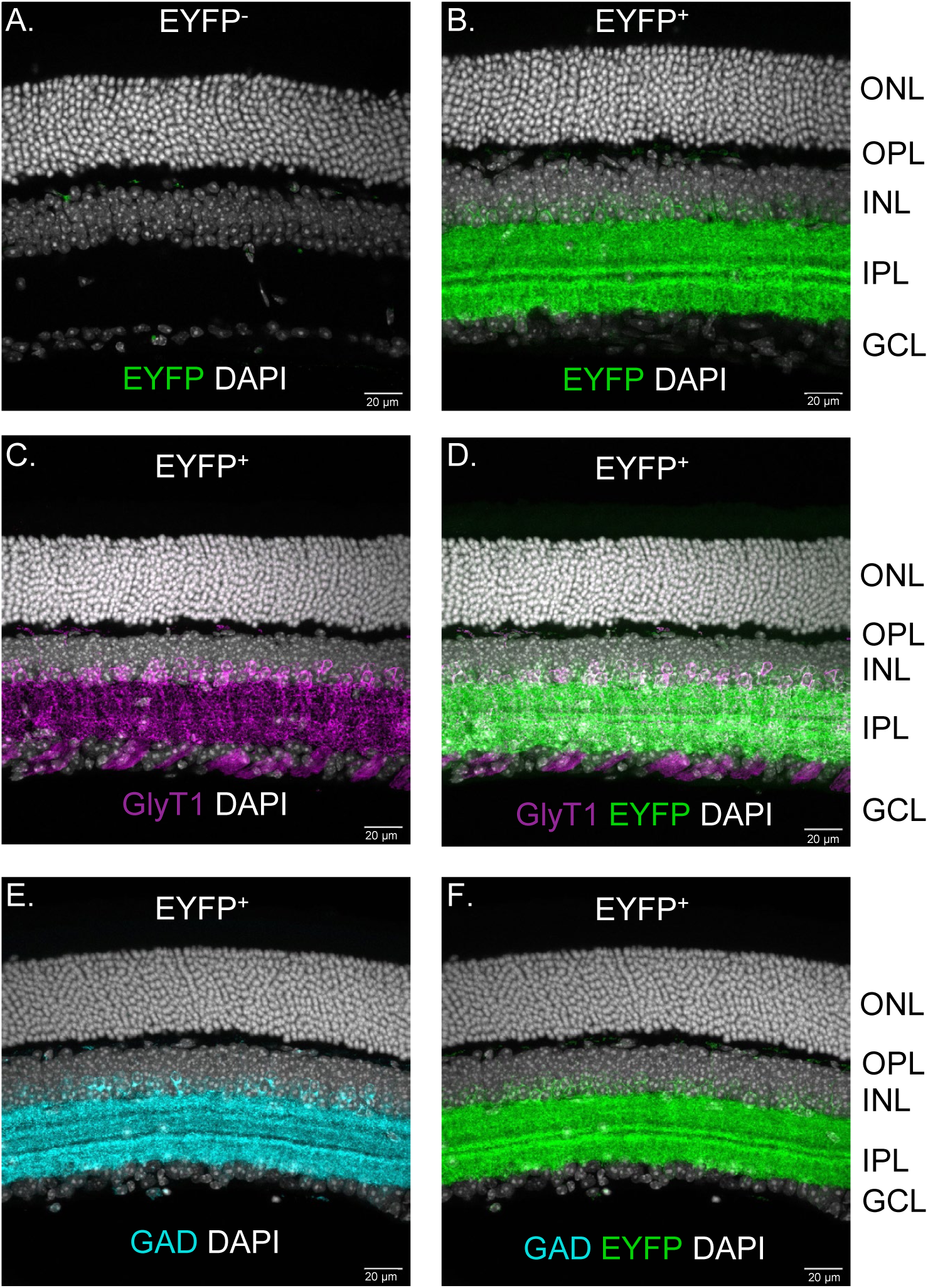
Retinal slices from VGAT-ChR2-EYFP^+^ mice selectively expressed EYFP in the OPL, INL and IPL. A. Retina from VGAT-ChR2-EYFP^-^ mouse does not show EYFP immunofluorescence (green). Nuclei labeled with DAPI (white). B. Retina from VGAT-ChR2-EYFP^+^ mouse shows EYFP immunofluorescence in the OPL, INL and IPL. C-D. Glycinergic cells (labeled for GlyT1, magenta) in VGAT-ChR2-EYFP^+^ retina overlapped with EYFP immunofluorescence (D). E. GABA releasing cells in VGAT-ChR2-EYFP^+^ labeled for GAD 65/57 (GAD, cyan) overlapped with EYFP immunofluorescence (F).

Whole-cell voltage clamp recordings from ACs while holding at -60 mV revealed fast oECs in response to 5 ms optogenetic light flashes (Fig. 2A). Mice not expressing EYFP did not have oECs in response to optogenetic light flashes (not shown). This confirms ChR2 is functionally expressed in the membranes of ACs. The amplitude and charge transfer (Q) of the oECs were light intensity dependent (Fig. 2B-C). The minimum light intensity necessary to evoke oECs was 1.18·10^9^ photons·µm^−2^·s^−1^. The maximum peak currents were measured at 8.35·10^9^ photons·µm^−2^·s^−1^ or maximum LED intensity. An intensity of 7.1·10^9^ photons·µm^−2^·s^−1^ was chosen for all subsequent stimuli to optogenetically evoke oECs and oIPSCs. The kinetics of the decay from the peak oECs were not light intensity dependent (Fig. 2D).

**Figure 2:**
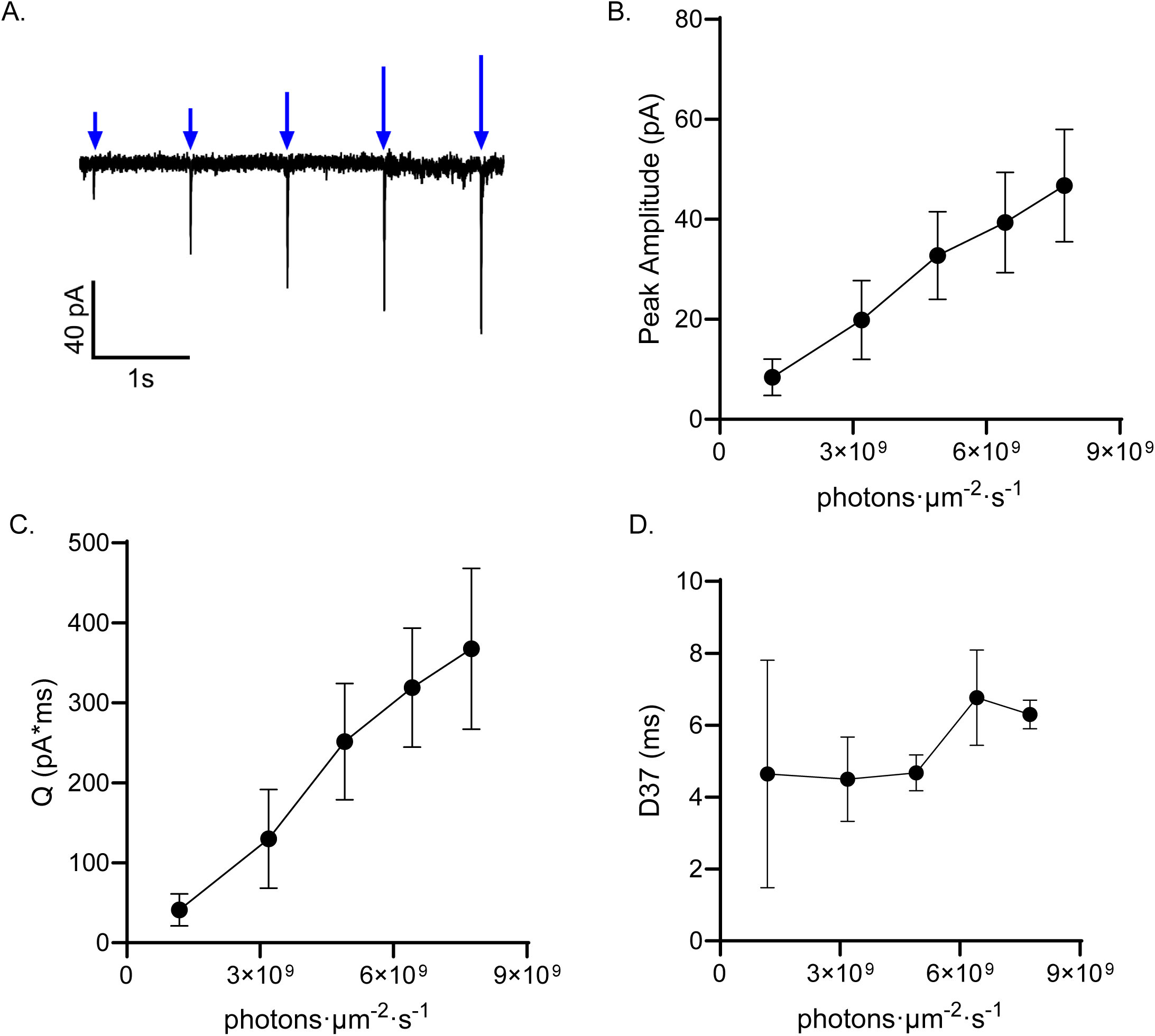
Optogenetic elicited excitation (oEC) of ChR2 expressing ACs is light intensity dependent. A. Example trace, oECs from ACs with 5 ms light stimuli at increasing intensities (values in B, stimuli = blue arrows). B-C. Peak amplitude (B) and Q (charge transfer, C) of oECs increases with increasing light intensity (p=0.01, n=5, RM-ANOVA). D. Decay time of oECs did not vary with intensity (p=0.66, n=5, RM-ANOVA). A minimum output of 1.18x109 photons·µm^−2^·s^−1^ was necessary to evoke an oEC.

In a physiologically light-evoked response, an OFF BC would only receive inhibitory inputs after photoreceptor activation followed by several synaptic pathways. Thus, the time to first peak and decay of IPSCs measured on OFF BCs is dependent on many presynaptic and post synaptic cell characteristics and activation which leads to slow activation and prolonged decay ^24^, potentially with a large contribution by Ca^2+^ mechanisms in ACs ^25^. To determine if AC GABA/glycine release shows a slow timecourse, even with brief optogenetic activation, we recorded AC oECs and oIPSCs during a full field 1 sec optogenetic light stimulus in VGAT-ChR2-EYFP^+^ (VGAT-ChR2) mice (Fig. 3A). In response to optogenetic stimulation, we observed significantly faster time to first peak and decay times of AC excitatory currents than would be observed in physiological light evoked responses ^26^ (Fig. 3B, C), presumably due to the fast opening and closing time of ChR2 channels ^27, 28^. However, the AC oIPSCs display kinetics similar to what would be expected for physiological light-evoked currents ^26, 29^. The fast opening and closing of ChR2 in oECs suggests the slower oIPSCs decays measured on postsynaptic cells were a function of synaptic kinetics and not from a prolonged depolarized or slow repolarizing effect in the presynaptic inhibitory neurons. This also suggests this technique is an effective way to isolate the AC to BC circuitry and determine if neuromodulators like dopamine can affect this circuit directly.

**Figure 3:**
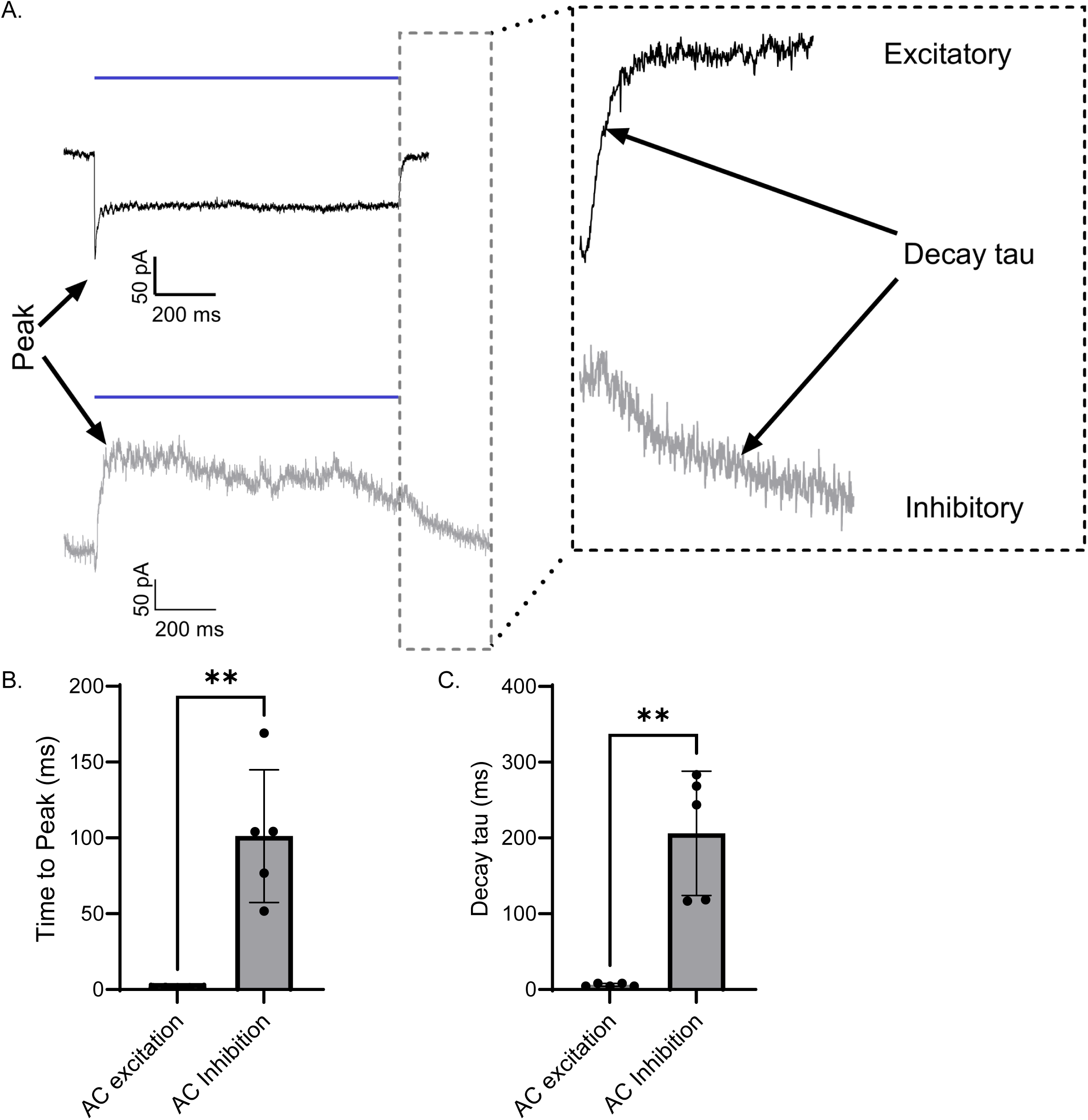
AC oECs and oIPSCs onto ACs have distinct kinetics. A. Example traces from the same AC: top oECs (V_m_ = -60 mV, blue line = 1 s light stimulus, 7.1·10^9^ photons·µm^−2^·s^−1^), bottom oIPSC (V_m_ = 0 mV). Left arrows show the peak locations. Inset shows magnified decay of current after light offset, right arrows show decay tau measurement. B. Rise time of oECs was significantly faster than that for oIPSCs from ACs (p=0.007, n=5, t-test). C. Decay tau of oECs was significantly faster than oIPSCs on ACs (p=0.006, n=5, paired t-test).

OFF BCs express GABA_A_Rs, GABA_c_Rs, and glycineRs. Direct activation of VGAT-ChR2 ACs elicited repeatable oIPSCs on OFF BCs (Fig. 4A). Optogenetic stimuli did not elicit oECs on OFF BCs (data not shown). There was also no difference in the peak amplitude, Q, or decay tau between the elicited inhibitory currents on OFF BCs before and after pharmacologically blocking PR inputs (data not shown). Both light adaptation and D1R activation with SKF reduce peak amplitude and Q of total inhibition onto OFF BCs in response to a 1 second physiological light stimulus ^13^. However, direct depolarization of ACs via 1 sec. optogenetic light stimulus elicited inhibition in OFF BCs that was not statistically different after application of SKF (Fig. 4A, p=0.43, t-test). OFF BCs expressed 2 distinct sets of currents. Some cells displayed many transient currents, which looked like individual spontaneous currents, during the 1 sec. light stimulus, and some displayed both transient and sustained currents. This suggests that optogenetic depolarization of ACs reveals unique release kinetics of certain neurotransmitters.

**Figure 4:**
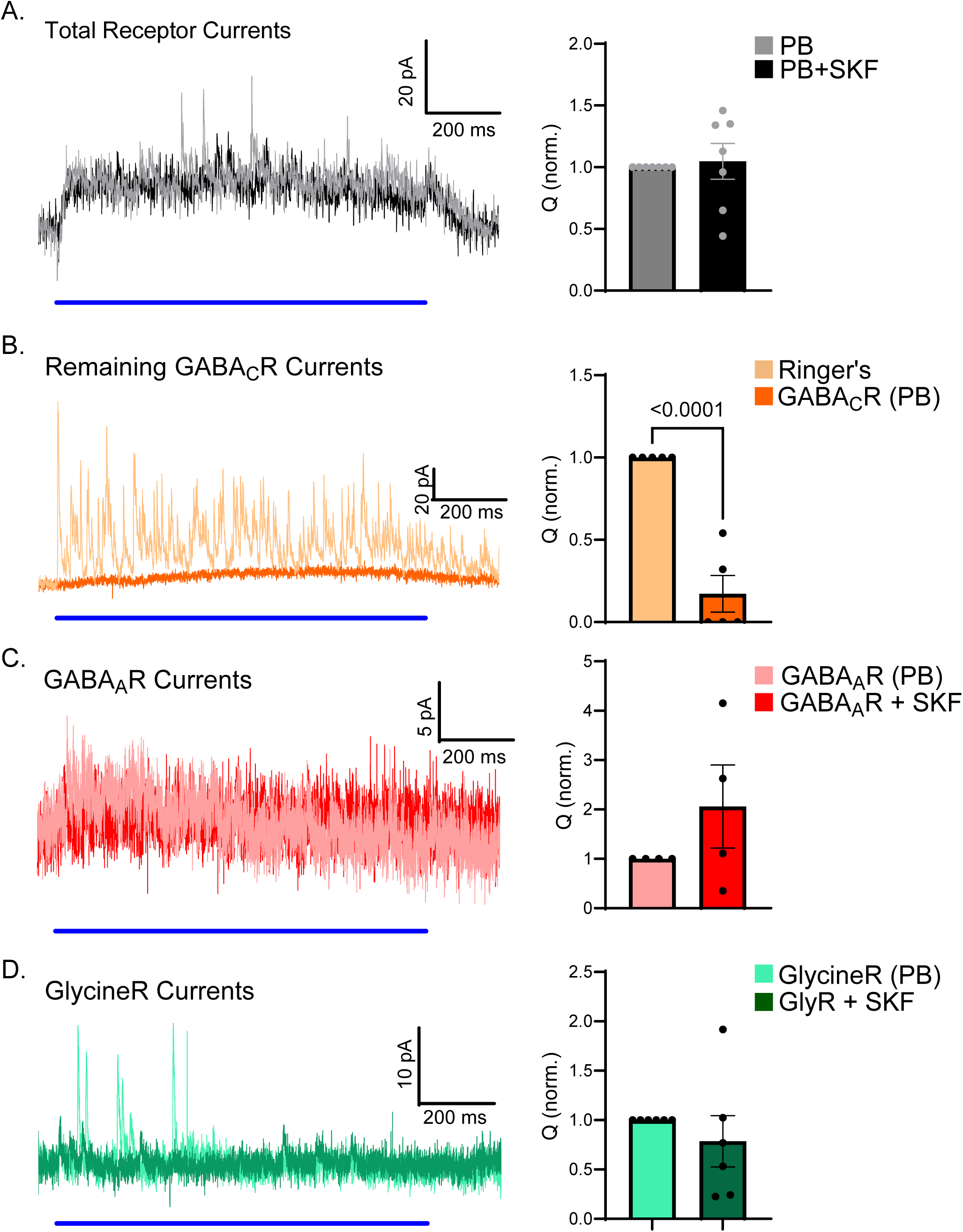
Components of oIPSCs from OFF-BCs. A. Example traces of total oIPSC (Left, in photoblock – PB, blue line = 1 s light stimulus, 7.1·10^9^ photons·µm^−2^·s^−1^) and oIPSC Q (right, normalized to control) before and after D1R agonist SKF was applied (p=0.43, n=6). B. Example of total oIPSC (left) and oIPSC Q (right, p<0.0001, n=5) before and after GABA_C_Rs were isolated (PB, strychnine, SR95531). C. Example GABA_A_R isolated oIPSC (left, PB, strychnine, TPMPA) and oIPSC Q (right, p=0.26, n=4) before and after D1R agonist SKF. D. Example glycineR isolated oIPSC (left, PB, SR95531, TPMPA) and oIPSC Q (right, p=0.43, n=6) before and after D1R agonist SKF.

To examine this, OFF BCs oIPSCs were recorded while isolating GABA_A_R, GABA_C_R, and glycineR mediated currents. Isolating GABA_C_R mediated currents with the addition of the photoblock cocktail (PB), GABA_A_R antagonist SR95531 (SR), and glycineR antagonist strychnine (see methods) eliminated all transient currents and severely reduced sustained oIPSCs (Fig. 4B). Q was reduced to 17.2 ± 11% of the total after GABA_C_R isolation (Fig. 4B, p<0.001, t-test). This suggests GABA_C_R inputs on OFF BCs mediate a small contribution to the total Q, consistent with previous reports showing GABA_C_R inputs make a small contribution to OFF BCs light-evoked IPSCs ^24^. GABA_A_R oIPSCs (PB, GABA_C_R antagonist TPMPA, strychnine) showed small sustained and transient currents only present during the optogenetic light flash. The peak amplitude (p=0.11, t-test) or Q (Fig. 4C, p=0.26, t-test) of the GABA_A_R oIPSCs were not significantly different after the application of SKF.

Previous data suggested that reductions in glycinergic inputs to OFF BCs contributed most of the light adapted and SKF induced reductions in inhibition ^13^. To examine this, glycineRs were isolated using TPMPA and SR. All cells displayed transient events after isolation of glycineRs. Surprisingly, there was no change in the total Q of the glycineR oIPSC after application of SKF (Fig. 4D, p=0.43, t-test). However, when analyzing the individual currents during the 1 sec. light stimulus (Fig. 5A), the peak amplitude (Fig. 5B, p<0.001, t-test), Q (Fig. 5C, p<0.01, t-test) and decay tau (Fig. 5D, p<0.01, t-test) of glycineR transient currents were significantly reduced after SKF. There was no change in the frequency of these transient events during the 1 sec. light stimulus (Fig. 5E, p=0.41, t-test).

**Figure 5:**
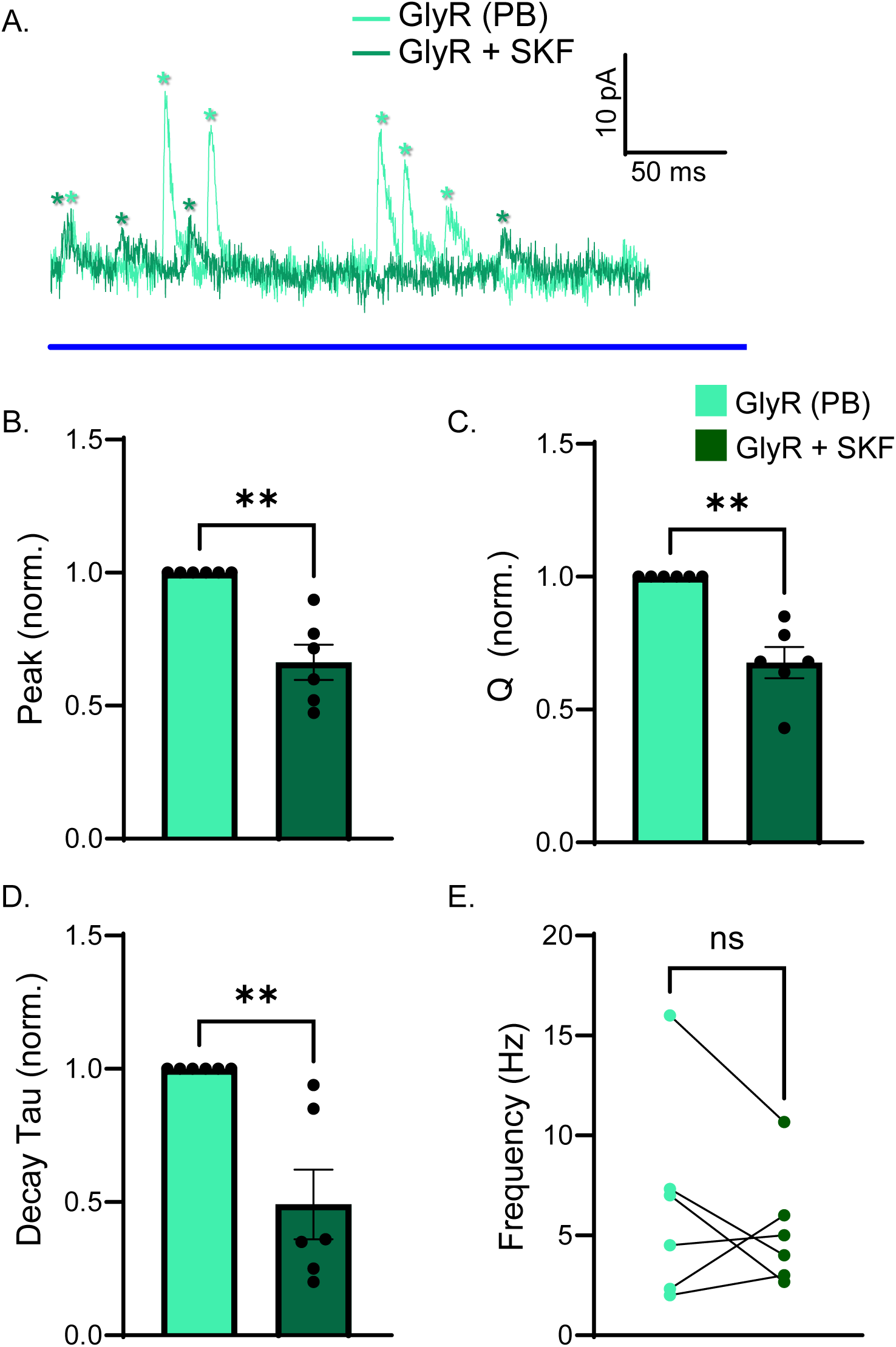
OFF-BCs glycineR transient currents during optogenetic stimulation are modulated by D1R activation. A. Transient currents analyzed during the light stimulus. Individual events marked with * (blue line = 1 s light stimulus, 7.1·10^9^ photons·µm^−2^·s^−1^). B. Average event peak amplitude was reduced after D1R agonist SKF application (SKF norm. to control, p<0.001, n=6 cells, t-test). C. Average event Q was significantly reduced by D1R activation (SKF norm. to control, p<0.01, n=6, t-test). D. Average event decay tau was significantly reduced by D1R activation (SKF norm. to control, p<0.01, n=6, t-test). E. Average event frequency was not changed by D1R activation (p=0.41, n=6, t-test). ** p<0.01

As shown in Fig. 3, presynaptic ACs also receive inhibitory inputs through release of GABA and glycine from other ACs onto GABA_A_Rs and glycineRs. This inhibition of inhibitory neurons is known as serial inhibition, which has been shown to modulate AC output onto BCs^30^. Serial inhibition is especially prominent when activating all inhibitory neurons by full field stimuli and could be affected by D1R activation. AC oIPSCs showed large, sustained currents with slow kinetics and also showed smaller transient currents (Fig. 6A). No differences were found in peak amplitude (Fig. 6B, p=0.29, t-test), Q (Fig. 6C, p=0.886, t-test) or time to first peak (p=0.314) of AC oIPSCs after the application of SKF. This suggests that the serial inhibition elicited as a result of AC activation does not play a role in observable SKF mediated changes in OFF BC oIPSCs.

**Figure 6:**
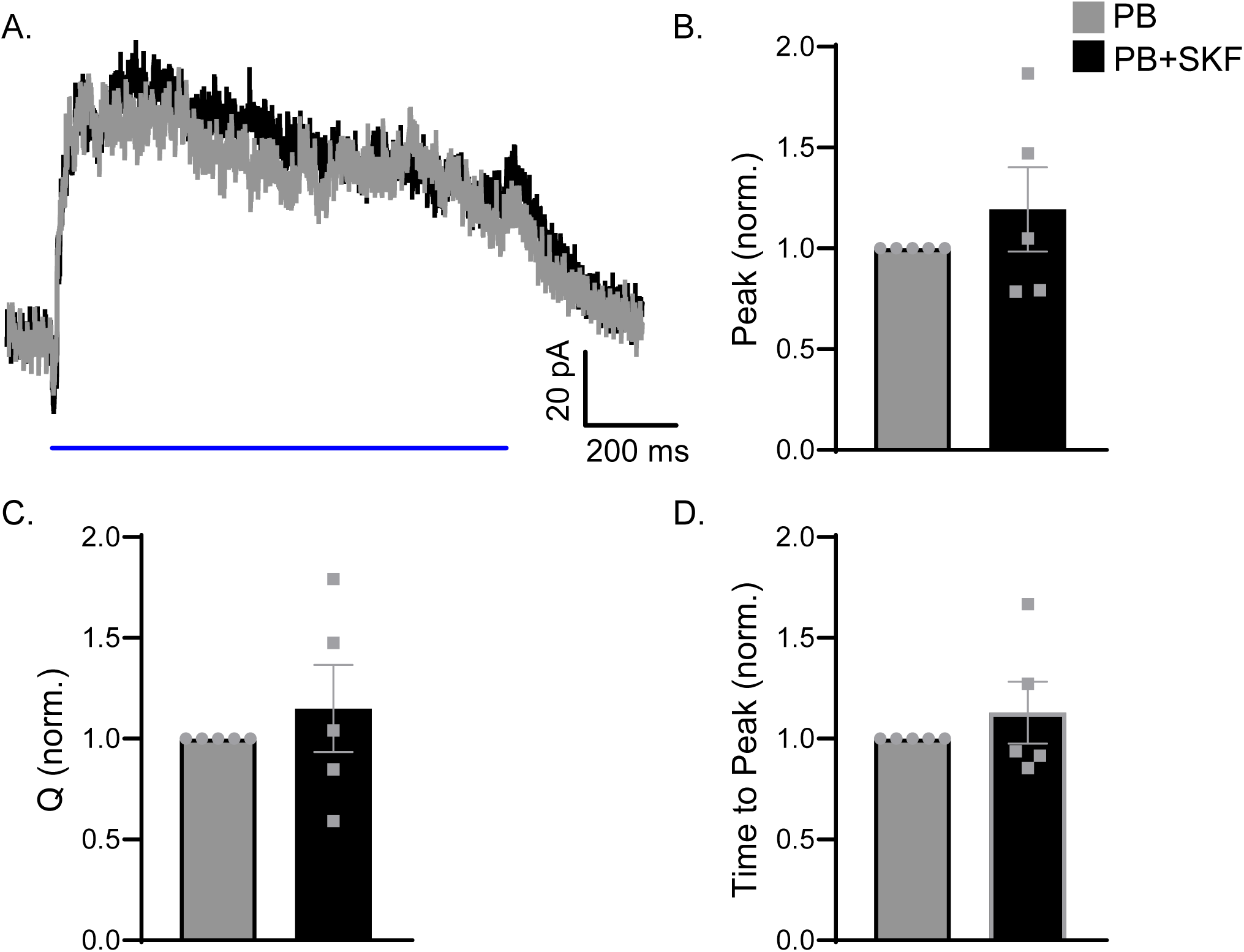
oIPSCs on ACs are not modulated by D1R activation. A. Example trace, oIPSCs from ACs (blue line = 1 sec stimulus, 7.1·10^9^ photons·µm^−2^·s^−1^, PB) with and without the D1R agonist SKF. B-D. The peak amplitude (B, p=0.29), time to first peak (C, p=0.886) and Q (D, p=0.314) of AC oIPSCs were unchanged after D1R application. SKF data is normalized to control, n=5 for all measurements.

## Discussion

Here we found that we could use the VGAT-ChR2 mice ^20^ to effectively activate isolated inhibition from ACs to OFF BCs that mimicked the timing of physiological light-evoked inhibition to OFF BCs. This slow timing of inhibition after ChR2 activation of ACs lends support to previous studies suggesting that neurotransmitter release from ACs is inherently slow and asynchronous ^25, 26, 31–35^ suggesting this could be a useful system to study neurotransmitter release kinetics. Using optogenetic control of GABA and glycine expressing cells, we found that the D1R agonist SKF reduced glycinergic inhibitory inputs onto OFF BCs that were elicited by direct activation of presynaptic ACs. Similar to previous findings with light-evoked inhibition ^13^, OFF BC oIPSCs are primarily glycinergic ^24^. Activation of D1Rs with SKF reduced glycinergic inhibition but not GABAergic inhibition, as had been previously shown with physiological light-evoked inhibition^13^. D1R activation has been shown to mimic reductions in inhibition on RBCs and OFF BCs due to light adaptation ^13, 14^. This suggests that increased dopamine release in light adapted retinas is driving modulations in synaptic communication. Since our current study directly activated ACs, our data suggest the AC to OFF BC synapse is modulated by dopamine independent of upstream dopaminergic effects previously studied ^8, 11, 36–40^.

In AC to OFF BC communication, dopamine mediated synaptic modulation could be expected to occur from the presynaptic AC and/or the postsynaptic OFF BC, because both neurons could express D1Rs ^17^. OFF BC glycinergic spontaneous inhibition frequency and amplitude have been shown to decrease in both light adapted conditions and after SKF application, which suggests at least some presynaptic effects ^13^. Though we observed reductions in peak amplitude and Q of glycinergic currents, we did not observe reductions in frequency of individual events after optogenetic stimuli. One explanation for the discrepancies between our data and previously measured light-evoked data ^13^ could be in the way ACs are being extensively depolarized. The strong, fast, and persistent inward cation influx through ChR2 could mask some presynaptic changes in release probability or amount of neurotransmitter seen in physiological light responses. Additional changes from D1R activation could come from upstream modulation of the ACs, either through their light activation from BCs or in reduced effects of serial inhibition between ACs ^30, 41, 42^, due to the ChR2-mediated extensive depolarization or the use of full-field stimuli instead of smaller bars that show spatial regulation of AC inhibition ^3^. If dopamine mediates these light-adapted changes in AC inputs, it is possible full field optogenetic stimuli do not capture these upstream changes. Mouse OFF BCs have been shown to express D1Rs differentially. Immunohistochemistry and in situ hybridization data have shown Type 1, 3b, and 4 OFF BCs exhibit D1R expression while type 2 and 3a do not ^15^. Morphological identification can be used to determine individual cell subtypes, but this is not a reliable technique for distinguishing between subtypes 1/2 or 3a/3b^21^. For this reason, our data have combined all OFF BC types we recorded from into our analysis. Identification of OFF BCs were confirmed by injecting Alexa Flour 568 during patch recordings. Since, type 2 and 3a OFF BCs do not express D1Rs, we would not expect these cells to be post-synaptically effected by SKF application. The potential incorporation of these cells into our data may limit the magnitude of our observed reduction in oIPSCs on OFF BCs if the role of dopamine in modulation of the OFF pathway is mediated by a post synaptic mechanism.

This work provides evidence that optogenetic control of specific neuron subtypes in the retina can be a useful tool in understanding neuromodulation in retinal circuitry. The vast locations of dopamine receptors in the retina give rise to many potential locations for synaptic control. The B6.Cg-Tg (Slc32al-COP4*H134R/EYFP) mouse line may also be a useful tool to study modulation of other inhibitory circuits in the retina as well.

